# Captivity affects mitochondrial aerobic respiration and carotenoid metabolism in the house finch (*Haemorhous mexicanus*)

**DOI:** 10.1101/2023.11.11.566700

**Authors:** Rebecca E. Koch, Chidimma Okegbe, Chidambaram Ramanathan, Xinyu Zhu, Ethan Hare, Matthew B. Toomey, Geoffrey E. Hill, Yufeng Zhang

## Abstract

In many species of animals, red carotenoid-based coloration is produced by metabolizing yellow dietary pigments, and this red ornamentation is an honest signal of individual quality. However, the physiological basis for associations between organism function and the metabolism of red ornamental carotenoids from yellow dietary carotenoids remains uncertain. A recent hypothesis posits that carotenoid metabolism depends on mitochondrial performance, with diminished red coloration resulting from altered mitochondrial aerobic respiration. To test for an association between mitochondrial respiration and red carotenoids, we held wild-caught, molting male house finches in either small bird cages or large flight cages to create environmental challenges during the period when red ornamental coloration is produced. We predicted that small cages would present a less favorable environment than large flight cages and that captivity would affect both mitochondrial performance and the abundance of red carotenoids. We found no evidence that living in small *versus* large cages had significant effects on wild-caught house finches; however, birds in cages of any size circulated fewer red carotenoids, showed increased mitochondrial respiratory rates, and had lower complex II respiratory control ratios—a metric associated with mitochondrial efficiency—compared to free-living birds. Moreover, among captive individuals, the birds that circulated the most red carotenoids had the highest mitochondrial respiratory control ratio for complex II substrates. These data support the hypothesis that the metabolism of red carotenoid pigments is linked to mitochondrial aerobic respiration in the house finch, but the mechanisms for this association remain to be established.

**SUMMARY STATEMENT:** Holding wild-caught male house finches in cages exposed a relationship between red carotenoid production and mitochondrial respiratory efficiency.

## INTRODUCTION

In many species of birds, the hue and chroma of carotenoid-based coloration is an honest signal of individual condition (Hill, 1991; Svensson and Wong, 2011). Numerous studies have documented associations between a host of proxies for the overall health and vigor (“condition”) of wild birds and the carotenoid-based coloration of both feathers and bare parts (Blount and McGraw, 2008; Hill, 2006). Experimental studies have demonstrated that carotenoid-based coloration can be sensitive to hormone manipulation (Khalil et al., 2020; McGraw et al., 2006a), dietary challenges (Hill, 2000; McGraw and Hill, 2000), infection by pathogens (Brawner et al., 2000; Hill et al., 2004; Thompson et al., 1997), and stress induced by captivity (Hill, 1992). The sensitivity of ornamental coloration to environmental stress has been found to be particularly striking for species that metabolically modify dietary carotenoids before they are used in ornaments (Brush, 1990; Mcgraw, 2006). Indeed, a meta-analysis investigating the strength of condition dependency of carotenoid-based ornaments in songbirds found that, relative to ornaments produced with dietary carotenoids, ornaments that required metabolized carotenoids were more reliable signals of condition (Weaver et al., 2018). Despite decades of study, however, the mechanisms linking the expression of carotenoid-based coloration to aspects of individual condition—and thereby the mechanisms by which such ornaments serve as honest signals—remain unclear.

Current hypotheses for the mechanisms by which carotenoid coloration can be a reliable signal of condition tend to fall into one of two categories: cost-based hypotheses that focus on the potential benefits of carotenoid pigments when they are not used as colorants, or index-based hypotheses that propose that shared pathways link expression of ornamental coloration to performance of physiological processes related to individual quality (Weaver et al., 2017). Testing these hypotheses requires an understanding of the mechanistic steps necessary to produce the colored ornament.

The shared pathway hypothesis proposes that display traits like red carotenoid coloration serve as reliable signals of condition because such traits indicate an individual’s capacity to maintain function of vital cellular processes in the face of environmental challenges (Hill 2011). In particular, the metabolism of red carotenoid pigments is proposed to be biochemically linked to aerobic respiration in mitochondria (the mitochondrial function hypothesis; Hill, 2014; Hill et al., 2019; Koch et al., 2017).

Various empirical observations support a link between mitochondrial parameters and production of red ketocarotenoids (which have a ketone group on one or both end rings; reviewed in Powers & Hill, 2021). A study of zebra finches (*Taeniopygia guttata*) found that treatment with mitoQ, a synthetic ubiquinone plus decyl- triphenylphosphonium (dTPP+) that acts as targeted mitochondrial antioxidant (Murphy and Smith, 2000; Murphy and Smith, 2007), increased bill redness, while treatment with only dTPP+ caused bills to become less colorful (Cantarero and Alonso-Alvarez, 2017). In a study of red crossbills (*Loxia curvirostra*), Cantarero *et al*. (2020) treated wild-caught males molting in captivity with either mitoQ or mitoTEMPO, a superoxide dismutase mimic (antioxidant) that is also targeted to the inner mitochondrial membrane. Interestingly, mitoQ treatment had no effect on the redness of feathers, though mitoTEMPO significantly increased the concentration of ketocarotenoids in plumage and hence increased plumage redness—but only among birds that were bright red at the time of capture (Cantarero et al., 2020). In other words, birds that appeared to be in the best condition at capture benefited from the effects of mitoTEMPO, but birds in lower condition did not.

These studies provide support for functional links between the production of ketocarotenoids and mitochondria, though questions remain as to the specific effects of the targeted mitochondrial manipulations and how we may extrapolate these findings to natural variation in wild systems. Perhaps the most direct test of links between mitochondrial aerobic respiration and ketocarotenoid production to date was a study of wild house finches (*Haemorhous mexicanus*), a species for which previous experimental studies have validated that males convert yellow dietary carotenoids to red ketocarotenoids that they use to color their feathers (Hill et al., 1994; Inouye et al., 2001; McGraw et al., 2006b; Toomey and McGraw, 2010). A study of molting male house finches found high concentrations of ketocarotenoids in the mitochondria of liver cells (Ge et al., 2015), supporting the idea that mitochondria play a role in carotenoid metabolism. A subsequent study found house finches molting in the wild showed positive associations between feather redness and various aspects of mitochondria, including inner membrane potential, respiratory control ratio (RCR), and the generation of new mitochondria (Hill et al., 2019).

Here, we compared the effects of environment perturbation during molt on both mitochondrial aerobic respiration and production of red feather pigments in male house finches. We placed wild-caught house finches into one of two cage sizes during the growth of feathers with ornamental coloration. Half of the captive birds were held in small bird cages in which they could hop but not fly, while the other half were held in large outdoor flight cages in which they could engage in short flights. We expected that birds would be subjected to a more challenging environment in small cages than in large cages, but that all captive birds would be in a more challenging environment than free-living birds. Several studies have shown that house finches develop drab yellow coloration when they molt in a cage, suggesting that the conditions of captivity compromise ketocarotenoid metabolism (Brush and Power, 1976; Grinnell, 1911; Hill, 1992; Hill, 2002).

We predicted that if captivity-associated loss of red coloration is associated with altered mitochondrial performance, then the challenge of living in a cage would affect both mitochondrial aerobic respiration and concentrations of circulating red ketocarotenoids, while concentrations of non-ketocarotenoids that are accumulated from the diet would not be affected. Maintaining birds under the controlled setting of captivity, and on a common diet, allowed us to experimentally test hypothesized relationships between mitochondrial respiratory performance and the production of red ketocarotenoids.

## METHODS

### Animals and housing

Wild male house finches at an early stage of molt were captured in Lee County, Alabama, USA between 1 and 5 August, 2022 following methods in Hill (2002). We estimated percent completion of molt by lifting body feathers to look for growing follicles beneath, and a percent of progression of molt of total body feathers was recorded. Birds had completed between 1 and 60% (with an average of 18% ± 3%, mean ± SE) of their prebasic molt, which is a complete or nearly complete replacement of feathers that spans about 100 days for most house finches (Hill 1993). This end-of-the-breeding-season molt is the only time that house finches can change the carotenoid-based pigmentation of their feathers, and hence, the key time point for evaluating production of red ketocarotenoids. At capture, we collected a sample of blood for analysis of circulating carotenoids in the plasma. We assigned birds to one of two age classes, based on plumage characteristics: hatching year (HY) for birds that were born in the year they were captured, or adult (A) for birds that had hatched in a previous calendar year. With the early stage of molt of most birds that we captured, feathers had not emerged from sheaths, and it was not possible to consistently quantify the color of incoming feathers; however, we did measure the carotenoid content of the developing feather follicles of a subset of experimental birds and found that plasma carotenoid levels are a good predictor of feathers’ content (see results). We therefore use circulating ketocarotenoid levels as our measure of red carotenoid metabolism in this experiment.

Captured experimental birds were transported to Auburn University (Auburn, AL, USA) and were assigned to a small or large cage by the flip of a coin. The large cages were outdoor cages were metal frames covered in hardware cloth 2.5 m x 2.5 m x 7.2 m with concrete floors that allowed birds to fly. The small cages were typical pet bird cages 0.5 m x 0.5 m x 0.5 m, arranged three cages high on racks in a temperature-controlled room with broad spectrum light; these cages allowed birds to hop from perch to perch but did not allow full flight. We anticipated that small cages would present birds with a less optimal environment for molt compared with large cages, based on previous observations (Hill 2002).

The base diet of all captive birds was Mazuri Mini Bird Diet (Mazuri Exotic Animal Nutrition, St. Louis, MO, USA), which is packaged as small pellets and formulated to provide complete nutrition for songbirds. We tested the carotenoid content of the formulated diet (see below) and per gram, it provided approximately 5 μg of lutein, 3 μg zeaxanthin, and 1 μg of β-carotene. We also added β-cryptoxanthin to the diets of birds by coating the pellets in organic papaya powder (Micro Ingredients, Montclair, CA, USA). Dried papaya is rich in β-cryptoxanthin, which has previously been hypothesized to be the main substrate that house finches and some other red cardueline finches use to produce their primary red pigment, 3-OH-echinenone (Hill, 2000; Stradi et al., 1997). Tests of the carotenoid content of pellets coated using our techniques recovered approximately 0.3 μg/g of pellets. Thus, lutein and zeaxanthin were the primary carotenoids available to birds, but small quantities of β-cryptoxanthin were also provided in diets.

For logistical reasons, all measurements of mitochondrial respiration had to be made in a four-day period, so birds were collected from the wild over four days and then processed in a four-day window, 14-15 days later. Specifically, on 16 to 19 August 2022, males were removed from cages and, after a blood sample was taken, were euthanized. These birds were dissected immediately, and mitochondria were isolated from their liver cells for analysis of respiration, as described below.

To provide broader context to the measurements of birds that had molted in cages, we also captured free-living males on 19 to 21 August 2022 and euthanized them at capture. For these males, we recorded age class and extent of molt (as above), and we took a sample of blood for carotenoid analysis. These birds were immediately dissected, and mitochondria were isolated from their liver cells for analysis of respiration as described below. The estimated percent of molt completed was comparable between the captive birds at the end of the experiment (80% ± 2%) and free-living birds (74% ± 2%).

All work with live animals was approved by the Auburn University Institutional Animal Care and Use Committee (2022-5048).

### Mitochondria isolation and respiration measurement

Mitochondria were isolated from the outer section of the right lobe of the liver according to Rogers *et al*. (2011) and Hill *et al*. (2019). Briefly, each liver sample was homogenized and subjected to differential centrifugations in isolation buffer (250 mM sucrose, 2 mM EDTA, 5 mM Tris-HCI, and 1% BSA, pH 7.4) on ice. Minced liver was first homogenized in a Potter-Elvehjem PTFE pestle and glass tube. The homogenate was centrifuged at 500 g for 10 minutes (4°C), then the supernatant was collected and centrifuged at 3500 g for 10 minutes (4°C). The resultant supernatant was discarded, and the final pellets (containing mitochondria) were suspended in ice-cold Mitochondrial Assay Solution (MAS-1: 2 mM HEPES, 10 mM KH_2_PO_4_, 1 mM EGTA, 70 mM sucrose, 220 mM mannitol, 5 mM MgCl_2_, 0.2% w/v fatty acid-free BSA, pH 7.4) and were kept at high concentration (∼20 mg protein/mL) on ice until use, according to Mookerjee *et al*. (2018). Total protein (mg/mL) was determined for each sample using Bradford assay reagent (Catalogue # 5000002, Bio-Rad, Hercules, California, USA). Liver mitochondria (0.175 mg/mL) respiration was measured in MAS-1 at 40°C using high resolution respirometry (Oroboros O2k, Innsbruck, Austria) according to Yap *et al*. (2022). For every sample, we measured respiration separately using either complex I (10 mM pyruvate, 10 mM glutamate, 2 mM malate) and complex II (10 mM succinate, 2 µM rotenone) substrates. For each complex, state 3_ADP_ respiration (hereafter, “state 3”) was induced by addition of 5 mM ADP, and state 4_O_ (state 4_Oligomycin_; hereafter, “state 4”) respiration was induced by addition of 2 µg/mL oligomycin. Non-mitochondrial respiration was induced by addition of 2.5 □M of antimycin A. Non-mitochondrial respiration was subtracted from state 3 and state 4 respiration before analysis. RCR was calculated by dividing state 3 by state 4 respiration. RCR can be interpreted as a proxy for mitochondrial “efficiency,” but it is more precisely defined as maximal capacity for respiration that results in ATP production relative to baseline respiration that offsets proton leak.

### Carotenoid analysis

We extracted and analyzed carotenoids from 5 or 10 μL of each plasma sample: 10 μL when possible, and 5 μL when the total sample volume was less that 10 μL. To each sample, we first added 250 μL of 100% ethanol and vortexed, then added 250 μL of hexane:*tert*-butyl methyl ether (1:1, vol:vol; hexane:MTBE), vortexed again, and centrifuged at 10,000 g for 3 minutes. We then transferred the supernatant (containing extracted carotenoids) to a separate 2 mL glass vial and evaporated it completely under a constant stream of nitrogen. For high performance liquid chromatography analysis (HPLC) of the extracted carotenoids, we dissolved each dried sample in 120 μL of mobile phase (acetonitrile:methanol:dichloromethane, 44:44:12, vol:vol:vol) and injected 100 μL into an Agilent 1200 series HPLC (Agilent, Santa Clara, CA, USA) with a YMC carotenoid column (5.0 μm, 4.6 mm × 250 mm; CT99S05-2546WT; YMC America, Inc., Devens, MA, USA) held at 30°C. We eluted samples with a mobile phase of acetonitrile:methanol:dichloromethane (44:44:12) for 11 minutes, which ramped up to acetonitrile:methanol:dichloromethane (35:35:30) from 11-21 minutes, then was held at isocratic conditions until 35 minutes; solvent was pumped at a constant rate of 1.2 mL/min throughout. We monitored sample elution using a UV-Vis photodiode array detector at wavelengths of 445 and 480 nm (for non-ketocarotenoids and ketocarotenoids, respectively), and we identified carotenoids through comparison to authentic standards (a gift of dsm-firmenich, Stroe, Netherlands) or to published accounts (Britton et al., 2004; Inouye et al., 2001; Potticary et al., 2020). We quantified each carotenoid peak by comparison to external standard curves of zeaxanthin for non- ketocarotenoids (detection limit 0.000203 μg) and astaxanthin for ketocarotenoids (detection limit 0.0003 μg), then calculated the concentration of that carotenoid in the plasma sample by adjusting for original sample volume (i.e. 5 or 10 μL), resuspension volume, and injection volume. We identified three major non-ketocarotenoids (lutein, zeaxanthin, and β-carotene) and two major ketocarotenoids (3-OH-echinenone and 4- oxo-rubixanthin) across our samples. For each captive individual, we obtained plasma carotenoid data from two time points: “pre-experiment” values from samples taken on initial capture, and “post-experiment” values from samples taken after captivity (at the same time as the mitochondrial measures). From the free-living birds, we obtained a single measurement of plasma carotenoid values from the same time point as when mitochondrial respiration was measured.

We also quantified carotenoid content from a sample of the papaya-coated pellet diet provided to the captive house finches. We followed a nearly identical extraction and measurement protocol as that described above, with a few exceptions. Pellets were first softened in 500 μL of 0.9% NaCl solution and then ground in Beadbug homogenizer (Benchmark Science, Inc., Sayreville, NJ, USA) with 0.1 g of zirconia beads (ZROB10; Next Advance, Inc., Troy, NY, USA) for 60 s at 4 kHz, before extracting carotenoids using ethanol and hexane:MTBE as described above. Then, after initial extraction and drying, we saponified the carotenoids (to hydrolyze carotenoid esters) by dissolving them for 6 hours in a 0.2 M solution of NaOH in methanol, and re-extracted using the same procedure as previously. We dissolved our final dried carotenoid extract in 120 μL of HPLC mobile phase, but we injected only 10 μL into the HPLC column for analysis.

Lastly, we measured carotenoids from the growing carotenoid-pigmented feather follicles of a subset of captive birds from the experiment. We obtained sufficient growing follicles from 19 males. We compared concentrations of carotenoids in feather follicles to concentrations circulating in plasma to validate the assumption that circulating 3-OH- echinenone levels are comparable to the levels deposited in the growing feathers. We collected an average of 3.0 mg (± 0.47 mg) per individual of whole follicles from frozen skin samples and extracted and analyzed carotenoids using an identical method as described above for the pellet diet, except we omitted the saponification step as a preliminary test revealed carotenoid esters to not be a major component of the follicle carotenoids.

### Statistical analyses

We performed all statistical analyses in R (v. 4.2.3; R Core Team, 2023) in RStudio (RStudio Team, 2023). First, we explored relationships among the different carotenoids measured using Pearson correlation matrices. We then focused our analyses on 3-OH- echinenone, the ketocarotenoid that is the largest component of both circulating carotenoids and ornamental coloration in the house finch (McGraw et al., 2006b). For all models, we first used a box-cox transformation on the response variable (one value of 0 μg/mL 3-OH-echinenone changed to the HPLC detection limit adjusted for sample volume to allow for lambda calculation), so the distribution of the data points did not differ significantly from normal (p > 0.05 in Shapiro-Wilk test).

To investigate potential precursor-product relationships among carotenoids, we first fit a simple linear model with transformed post-experiment 3-OH-echinenone as the response variable, and pre-experiment lutein and zeaxanthin (potential dietary precursor carotenoids), pre-experiment 3-OH-echinenone concentration (to control for variation in starting values), molt percent (score of 0-100), cage size treatment (small or large), and age class (A or HY) as fixed effects. We also fit a nearly identical linear model to test the relationship between normalized 3-OH-echineone detected in growing feather follicles and circulating 3-OH-echinenone, also including molt percent, cage size treatment, and age class as fixed effects.

To test for effects of cage size treatment during captivity on carotenoid levels, we fit linear models with either transformed post-experiment total non-ketocarotenoids or 3- OH-echinenone as the response variables, and cage size treatment, age class, molt percent, and total pre-experiment non-ketocarotenoids or 3-OH-echinenone (respectively) as fixed effects.

Due to the demands of running a high volume of samples in a low-throughput and time-sensitive process for evaluating mitochondrial respiration, we took a conservative approach in first removing statistically significant outlier measurements that may represent technical errors in our dataset of mitochondrial measurements. We tested for outliers separately in our measures of state 3 respiration, state 4 respiration, and RCR for complex I and complex II using the Grubbs test in the “outliers” package (v. 0.15; Grubbs, 1969; Komsta, 2022). We removed any data points that were statistically significant outliers, and we also removed any RCR value that was associated with a state 3 or state 4 measure that itself was an outlier. In total, we detected and removed two outliers from complex I state 4 and the corresponding two from complex I RCR, one from complex II state 3, one from complex II state 4, and four from complex II RCR (including the two corresponding to the state 3 or 4 outliers).

Next, we explored relationships among our mitochondrial respiration measures using Pearson correlations, as above. We found moderately high correlations between state 3 and state 4 measures within a complex (0.7-0.8, see below; Figure S1). While both measures are biologically distinct in terms of the aspect of mitochondrial respiration they represent, it is not unexpected that individual samples tended to have either higher or lower overall respiration rates within a complex, creating covariation between measures. We therefore include state 3 respiration in our main models (but exclude state 4) as a fixed effect to account for variation in respiration rate without introducing problematic collinearity. Indeed, we calculated variance inflation factors (VIFs) to gauge collinearity among the fixed effects for our models of mitochondrial respiration data using the “car” package (v. 3.1.2; Fox & Weisberg, 2019), and all VIFs < 2 (most < 1.5), suggesting that collinearity is not playing a major role in our effect estimates. In comparison, versions of these models run with both state 3 and state 4 measures included as fixed effects had VIFs of > 5.

To test whether variation in mitochondrial respiratory measures might predict variation in circulating 3-OH-echinenone levels at the end of the experiment, we fit two linear models, one for complex I and the other for complex II. Each model comprised normalized post-experiment 3-OH-echinenone as the response variable, and fixed effects of the respective complex’s state 3 respiration and RCR measures, cage size treatment, molt percent, and age class. Then, to test whether mitochondrial respiration measures might predict the magnitude of change in circulating 3-OH-echinenone between the start and end of the experiment (i.e. decrease after time in captivity), we first we calculated the percent loss of 3-OH-echinenone concentration relative to the starting concentration (i.e. starting concentration – ending concentration / starting concentration). To evaluate a percentage as a response variable, we fit linear models on data with an angular transformation (arcsin-square-root) applied. We again fit two models—one for each respiratory complex measured—with transformed percent 3-OH- echinenone lost as the response variable, and fixed effects of state 3 respiration, RCR, cage size treatment, age class, and molt percent.

We also we compared the measurements of free-living birds to those of our captive birds. We fit linear models for each measurement of interest, containing a box-cox transformed response variable (3-OH-echinenone levels, total non-ketocarotenoid levels, and state 3, state 4, or RCR for each of complex I and complex II) and fixed effects of captivity (captive *versus* free-living), molt percent, and age class. These comparisons helped us better capture the effects of captivity on mitochondrial respiration measures, given that—unlike circulating carotenoid levels—we could only quantify mitochondrial performance in each individual once.

## RESULTS

### Relationships among carotenoids

HPLC analyses of plasma samples revealed that both wild and captive-held birds had carotenoid types and concentrations typical of house finches (McGraw et al 2006). As expected among molting birds, the ketocarotenoid 3-OH-echinenone was the most abundant carotenoid in plasma overall, followed by the ketocarotenoid 4-oxo- rubixanthin, and then the non-ketocarotenoids lutein and zeaxanthin. We also detected scant amounts of β-carotene in some samples, but notably, no measurable β- cryptoxanthin—previously implicated as a main 3-OH-echinenone precursor (McGraw et al 2006)—was detected in any plasma sample (either in free-living birds or captive-held birds).

Captive birds at the end of the experiment had lower circulating 3-OH- echinenone levels than free-living birds (p < 0.001), but captive and free-living individuals did not differ in total non-ketocarotenoid levels (p = 0.89; Figure 1; Table S1), suggesting that being held in captivity had a negative effect on ketocarotenoid production but not absorption and circulation of dietary carotenoids. Interestingly, cage size did not affect circulating 3-OH-echinenone, though birds held in small cages had significantly more circulating non-ketocarotenoids than birds in large cages (Figure 1; Table 1). In general, captive birds decreased circulating ketocarotenoids and non- ketocarotenoids alike between the start and end of the experiment (ketocarotenoids: 48.0 ± 3.9 μg/mL before experiment, 13.1 ± μg/mL after experiment; non- ketocarotenoids: 12.3 ± 0.8 μg/mL before experiment, 9.2 ± 0.9 μg/mL after experiment; mean ± SE). We found that post-experimental levels of circulating 3-OH-echinenone were strongly predicted only by pre-experimental levels of that carotenoid (p < 0.001), indicating that the same birds with high initial circulating concentrations of 3-OH- echinenone tended to have high final concentrations; we did not detect any effect of pre-experimental non-ketocarotenoid levels (lutein and zeaxanthin) on post-experiment 3-OH-echinenone (p > 0.2; Table S2), suggesting that birds were not limited in production of this ketocarotenoid by the availability of these two dietary carotenoids.

**Figure 1.**
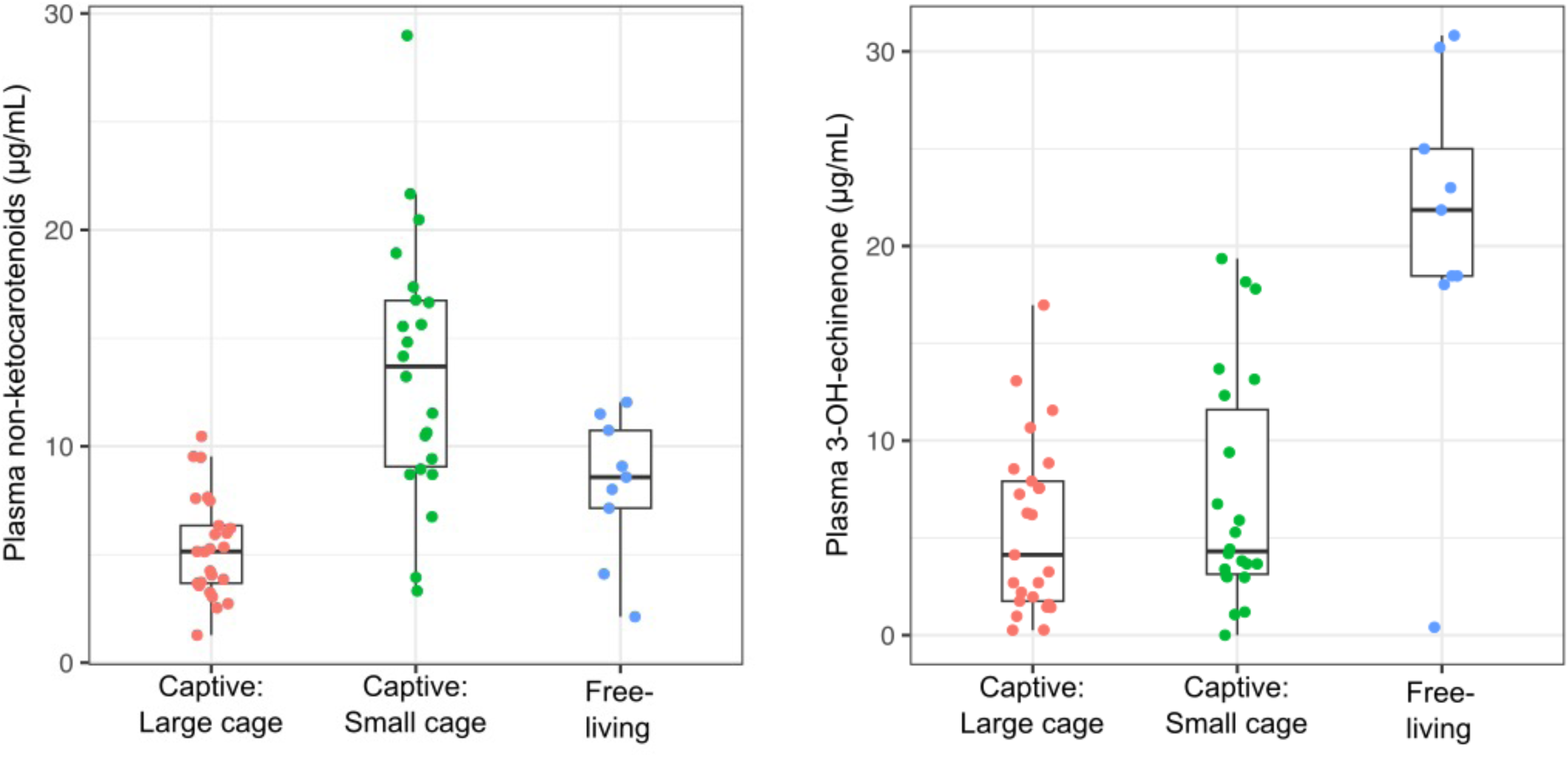
The concentrations of non-ketocarotenoids or 3-OH-echinenone in the plasma of house finches that were either held in small bird cages, housed in large flight cages, or free-living. All birds were growing feathers with carotenoid pigments at the time measurements were taken. Points are measurements from each bird. Box plots show median, interquartile range, and confidence limits of sample.

**Table 1.** Results of linear models testing the effects of cage size treatment, age, molt percent, and initial (pre-experimental) levels of circulating non-ketocarotenoids (top) or 3-OH-echinenone (bottom) on levels in circulation at the end of the experiment.

|  | Effect | Estimate | Std. Error | *t* | *P* |
| --- | --- | --- | --- | --- | --- |
| Non-ketocarotenoids | Intercept | 1.79 | 0.94 | 1.90 | 0.06 |
|  | Pre-experiment levels | 0.02 | 0.02 | 1.34 | 0.19 |
|  | Molt percent | -0.0030 | 0.01 | -0.31 | 0.76 |
|  | Treatment (small) | 1.12 | 0.18 | 6.23 | <0.001 |
|  | Age (hatch year) | -0.13 | 0.25 | -0.53 | 0.60 |
| 3-OH-echinenone | Intercept | -1.80 | 1.81 | -0.99 | 0.33 |
|  | Pre-experiment levels | 0.09 | 0.01 | 6.32 | <0.001 |
|  | Molt percent | 0.02 | 0.02 | 1.18 | 0.25 |
|  | Treatment (small) | 0.45 | 0.36 | 1.24 | 0.22 |
|  | Age (hatch year) | -0.39 | 0.51 | -0.76 | 0.45 |

When applicable, the reference group for a categorical variable is listed in parentheses.

Among the males for which we could analyze carotenoids in growing feather follicles, circulating 3-OH-echinenone predicted the concentration of 3-OH-echinenone in the growing feather follicles (p = 0.001; Figure S2; Table S3), supporting our assumption that the concentration of this ketocarotenoid in circulation is a useful predictor of its levels in growing colored feathers in the captive birds. Interestingly, hatch-year birds deposited a higher concentration 3-OH-echinenone into their follicles relative to the amount circulating compared to adult birds (p = 0.037; Figure S2; Table S2), perhaps indicating that young birds adapt to capivity better than older birds. A larger sample size of adult birds would be necessary to probe this relationship further.

### Mitochondrial respiration measurements

When we compared mitochondrial respiration between captive and free-living birds, we found that the birds in captivity tended to have higher respiration measures than their free-living counterparts (complex I states 3-4 and complex II state 4, p < 0.04; complex II state 3 not significantly different, p = 0.12; Figure 2; Table S1). Captive birds also had lower complex II RCR than free-living birds (p = 0.026; Figure 2; Table S1). These results suggest that captivity largely increased mitochondrial respiration rates and also changed the ratio between state 3 and state 4 rates in complex II, causing decreased RCR.

**Figure 2.**
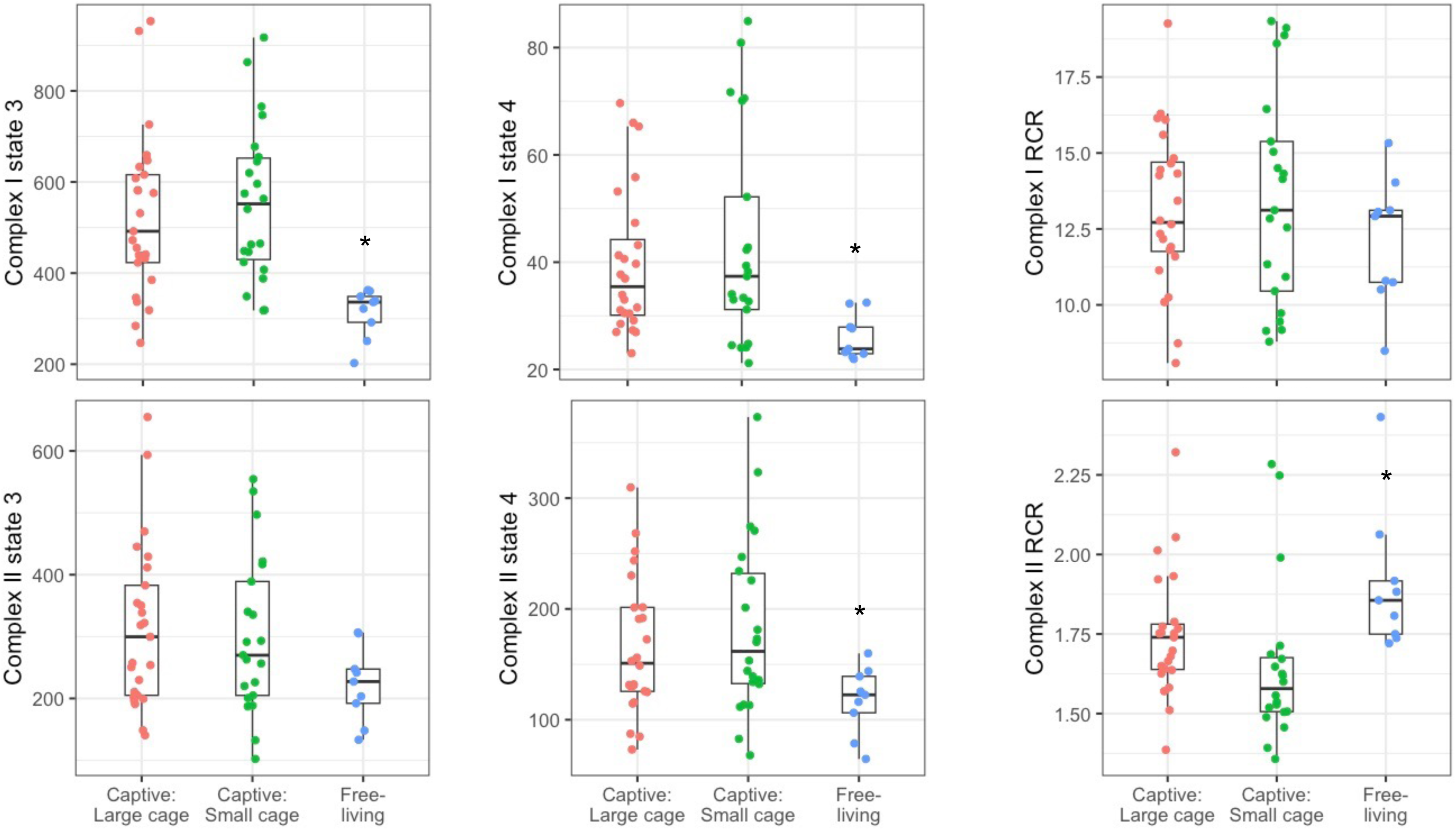
Measures of aerobic respiration in the mitochondria of liver cells of house finches that were either held in small bird cages, housed in large flight cages, or free-living. Complex I and complex II measurements were made by providing substrates that introduce electrons through complex I or complex II of the electron transport system (ETS), respectively. Respiratory control ratio (RCR) is derived by dividing state 3 by state 4 respiration. The units for state 3 and state 4 respiration are nmol O_2_ per min per mg protein. Asterisks indicate statistically significant differences between captive and free-living populations (Table S1). Boxplots described in Fig. 1.

We also found that captive house finches with higher complex II RCR circulated higher levels of 3-OH-echinenone (p = 0.029; Figure 3; Table 2). Captivity caused a significant reduction in average levels of circulating 3-OH-echinenone (Figure 1), and we were curious to determine whether mitochondrial respiration measures had an association with the magnitude of this decline. Therefore, we investigated the relationship between transformed percent loss of 3-OH-echinenone in captive birds between the start and end of the experiment, and mitochondrial respiration measures. We found that birds with higher complex II RCR maintained more circulating 3-OH- echinenone from the start to the end of the experiment (p = 0.042; Table 3). These results both suggest a link between complex II respiration and increased circulation of 3-OH-echinenone, while under the physiological challenge of captivity.

**Figure 3.**
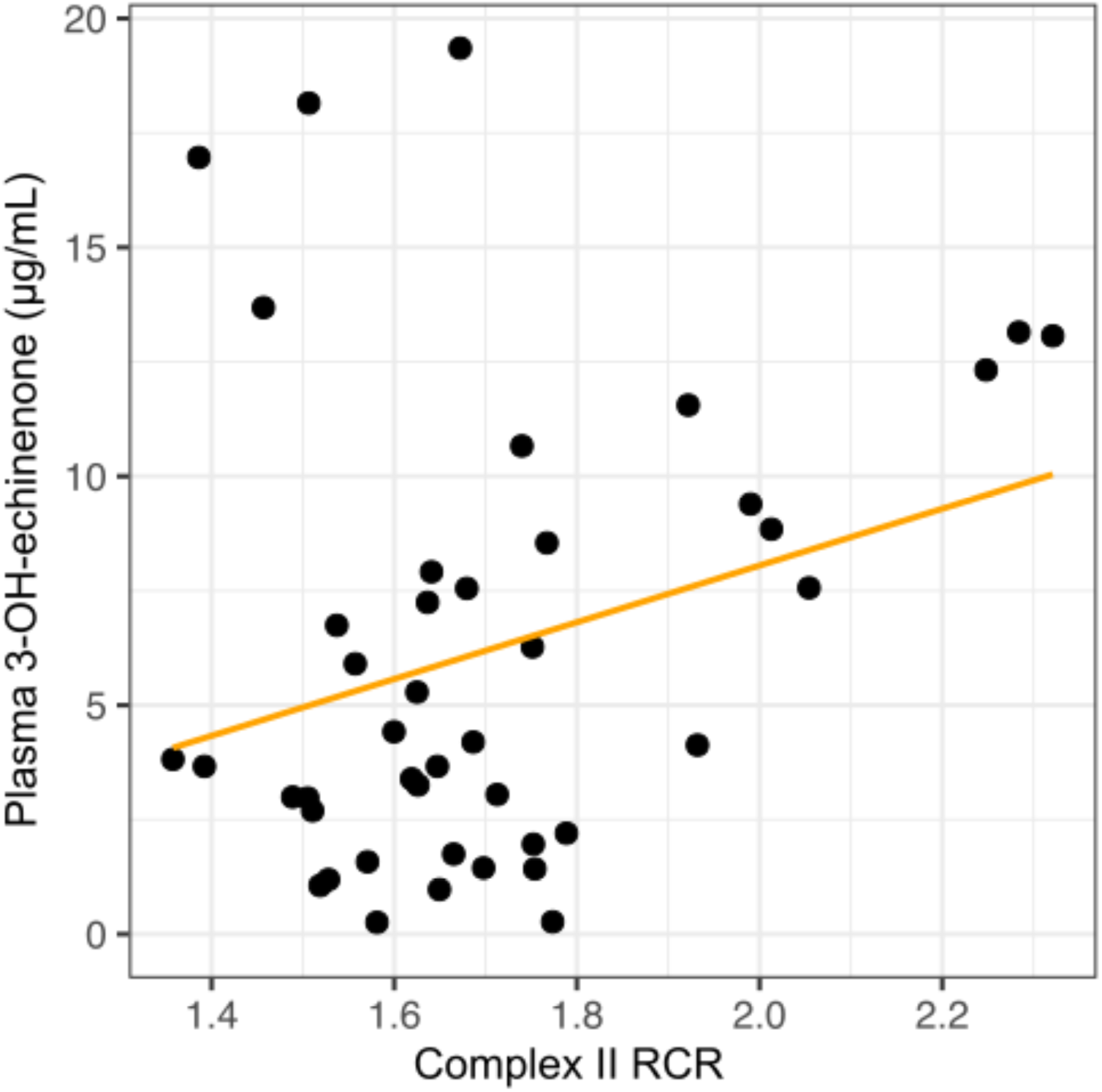
Scatterplot of the relationship between complex II RCR and circulating 3-OH-echinenone for captive-held birds. See Fig. 2 for definitions of mitochondrial measures.

**Table 2.** Results of linear models testing the effects of cage size treatment, age, molt percent, and complex I (top) or complex II (bottom) mitochondrial respiration measurements on circulating 3-OH-echinenone in captive birds at the end of the experiment.

|  | Effect | Estimate | Std. Error | *t* | *P* |
| --- | --- | --- | --- | --- | --- |
| Complex I | Intercept | -0.68 | 2.71 | -0.25 | 0.80 |
|  | Cage size (small) | 0.61 | 0.48 | 1.25 | 0.22 |
|  | Age (hatch year) | -0.14 | 0.68 | -0.2 | 0.84 |
|  | RCR | -0.14 | 0.081 | -1.76 | 0.087 |
|  | State 3 respiration | 0.0022 | 0.0016 | 1.38 | 0.18 |
|  | Molt percent | 0.042 | 0.026 | 1.60 | 0.12 |
| Complex II | Intercept | -4.11 | 2.80 | -1.47 | 0.15 |
|  | Cage size (small) | 0.69 | 0.46 | 1.48 | 0.15 |
|  | Age (hatch year) | -0.53 | 0.64 | -0.83 | 0.41 |
|  | RCR | 2.36 | 1.04 | 2.27 | 0.029 |
|  | State 3 respiration | 0.0034 | 0.0022 | 1.55 | 0.13 |
|  | Molt percent | 0.016 | 0.025 | 0.65 | 0.52 |

When applicable, the reference group for a categorical variable is listed in parentheses.

**Table 3.** Results of linear models testing the effects of cage size treatment, age, molt percent, and complex I (top) or complex II (bottom) mitochondrial respiration measurements on the angular-transformed percent loss of circulating 3-OH-echinenone in captive birds at the end of the experiment.

|  | Effect | Estimate | Std. Error | *t* | *P* |
| --- | --- | --- | --- | --- | --- |
| Complex I | Intercept | 1.25 | 0.38 | 3.28 | 0.002 |
|  | Cage size (small) | -0.11 | 0.068 | -1.61 | 0.12 |
|  | Age (hatch year) | 0.071 | 0.095 | 0.75 | 0.46 |
|  | RCR | 0.006 | 0.011 | 0.51 | 0.62 |
|  | State 3 respiration | -0.00005 | 0.00002 | -0.24 | 0.81 |
|  | Molt percent | -0.003 | 0.004 | -0.86 | 0.40 |
| Complex II | Intercept | 1.63 | 0.40 | 4.0 | <0.001 |
|  | Cage size (small) | -0.15 | 0.066 | -2.18 | 0.036 |
|  | Age (hatch year) | 0.083 | 0.091 | 0.911 | 0.368 |
|  | RCR | -0.313 | 0.149 | -2.104 | 0.042 |
|  | State 3 respiration | 0.000 | 0.000 | 0.134 | 0.894 |
|  | Molt percent | -0.001 | 0.004 | -0.207 | 0.837 |

When applicable, the reference group for a categorical variable is listed in parentheses.

## DISCUSSION

In this study, we explored a version of the shared pathway hypothesis for honest signaling that proposes that the efficiency of conversion of red ketocarotenoids from yellow dietary carotenoids depends on the efficiency of mitochondrial aerobic respiration—the mitochondrial function hypothesis (Cantarero et al., 2020; Hill, 2011; Hill, 2014; Powers and Hill, 2021). A previous study testing this hypothesis in free-living house finches found a statistical association between the redness of growing feathers and various measures of liver mitochondrial performance (Hill et al. 2019). Our goal in the current study was to further investigate links between mitochondrial respiration and the conversion of dietary carotenoids to ornamental red ketocarotenoids by experimentally altering cellular conditions through an environmental challenge during molt in a group of male house finches. We predicted that males held in small cages would be subjected to greater captivity effects in both mitochondrial measures and ketocarotenoid levels than males held in large outdoor flight cages, and that captivity would both alter mitochondrial aerobic respiration and depress circulating ketocarotenoid levels.

We observed four key outcomes of our experiment: 1) Captivity increased most rates of mitochondrial respiration while decreasing complex II RCR. 2) Captivity decreased circulating 3-OH-echinenone, the major red ketocarotenoid in house finches. 3) Among captive birds, males that circulated the most 3-OH-echinenone had the highest complex II RCR. And, 4) living in small, indoor cages had no greater impact on wild-caught house finches than did living in large, outdoor cages. Collectively, these findings suggest that holding house finches in captivity created an altered physiological state that both perturbed mitochondrial aerobic respiration and reduced the production of red ketocarotenoids like 3-OH-echinenone. Further, the association between higher circulating ketocarotenoid levels and higher mitochondrial complex II RCR revealed in captive birds supports the hypothesis that production of ketocarotenoids in house finches is responsive to aerobic respiration in the mitochondrion.

To our knowledge, the effects of captivity on mitochondrial aerobic respiration in birds have not been previously reported, though inferences can be drawn based on studies of related measurements. For example, a study of wild great tits (*Parus major*) found that administration of glucocorticoids—which can be broadly considered a treatment to increase physiological stress—caused increased mitochondrial proton leak, which is related to the state 4 respiration measured in our study (Casagrande *et al*., 2020). State 4 respiration—also referred to as baseline respiration—is measured under conditions where mitochondria are provided no ADP for production of ATP, so oxygen consumption comes from proton leakage across the inner mitochondrial membrane.

Such leak-related respiration has previously been hypothesized to be related to basal metabolic rate (BMR; Brand, 1990; Jastroch *et al*., 2010; Metcalfe *et al*., 2023), and short-term captivity in birds has previously been found to induce increased measures of BMR, though mitochondrial respiration was not measured in these studies (McNab, 2009; Weathers et al., 1983). In another study, treatment with a chemical that experimentally induces increased mitochondrial proton leak—a mitochondrial “uncoupler” (DNP, 2,4-dinitrophenol)—increased BMR in birds (Stier et al., 2014).

These lines of evidence suggest that increasing stress in wild birds by holding them in captivity may cause increased rates in whole-animal basal respiration as well as state 4 mitochondrial respiration, and that this increase may be driven by higher mitochondrial proton leak. Our findings in the current study are supportive of this effect of captivity: we found that house finches held in cages had higher state 4 measurements than free-living birds for tests of both complex I and complex II.

Interestingly, RCR significantly differed between captive and free-living house finches only for complex II; this may be due to significant increases in both state 3 and state 4 (leading to no significant change to their ratio) in complex I, but a significant increase in only state 4 in complex II (leading to a significant decrease in the ratio).

Such a pattern requires further testing, but it does suggest that complex II RCR is particularly sensitive to the conditions of captivity. This result is in line with our observation that complex II RCR, but not that of complex I, relates to concentration of ketocarotenoids in captive birds.

The observations in this current study corroborate those of a previous field study of house finches that found positive associations between the redness of plumage coloration and measures of mitochondrial respiration in wild molting males (Hill et al. 2019); however, the two studies differ in the mitochondrial parameters that were found to be associated with production of red pigments. In the current study, we observed significant associations between circulating ketocarotenoids and the RCR of complex II, while the previous study found an association between feather redness and the RCR of complex I (Hill et al. 2019). Given that house finch feather hue derives from the carotenoid pigments deposited while that feather is growing (Butler et al., 2011; Inouye et al., 2001), and that we detected a tight correlation between circulating ketocarotenoids and ketocarotenoids deposited in growing feathers in our experiment, we expect that the difference in results between the two studies is not likely to be due to measuring circulating ketocarotenoids *versus* feather hue. Instead, we consider that the different patterns may arise from differences between the substrates used by complex I and complex II. Complex I receives electrons from NADH, which is produced during the breakdown of glucose and carbohydrates during glycolysis as well during the citric acid cycle. In contrast, complex II receives electrons from succinate (FADH_2_), an intermediate in the citric acid cycle (Cooper and Adams, 2023). Interestingly, the oxidation of fat produces a higher percentage of FADH_2_ than NADH compared to using carbohydrates as substrates (Cooper and Adams, 2023). Consequently, the differences between the results of the two studies could have arisen from multiple sources: complex II respiration may be more sensitive to the type of environmental stress caused by captivity relative to complex I, or the altered diet of captive birds may have changed the relative amounts of carbohydrate and fat substrates available for respiration.

It is important to note that variation in mitochondrial aerobic respiration measures cannot be easily simplified down to “better” or “worse” performance, as mitochondria are dynamic and changes to aspects of cellular respiration, like amount of proton leak, can be flexibly adjusted to respond to current conditions (Koch et al., 2021; Monzel et al., 2023). Within the context of our current study, comparing results from our captive birds to measurements from free-living birds aids in interpreting the patterns we detected within the captive group. Given that we found captivity to cause decreased complex II RCR, we might expect that the individuals maintaining the highest complex II RCR despite captivity are those that are least perturbed by the change in environmental conditions. Following this logic, we might consider these individuals to be our highest “quality” birds, since they appear able to withstand the same challenge of being held captive while altering less of their cellular physiology than other individuals. That these birds also circulated more ketocarotenoids in captivity aligns with this perspective, and with the fact that redness in house finches has historically been found to correlate with other measures of quality. We therefore propose that the conditions posed by captivity revealed underlying variation in individual quality that was not detectable in free-living birds.

In conclusion, we found that confining birds to cages affected both mitochondrial respiration and the concentration of circulating red ketocarotenoids. Perhaps most significantly, captive individuals that maintained mitochondrial performance closest to that of free-living birds despite also produced the most ketocarotenoids. These observations support the hypothesis that carotenoid metabolism is linked to mitochondrial aerobic respiration. The specific mechanisms that that underlie an association between carotenoid ornamentation and mitochondrial function remain unclear, however, and more targeted experiments using chemical or genetic manipulations of specific components of both cellular respiration and carotenoid metabolism will be needed to further advance this field of study.

## Acknowledgements

University of Tulsa undergraduate students Hannah Reeb and Brooke Joski assisted with carotenoid extraction and analyses. Jennifer Parsons at Mazuri Exotic Animal Nutrition provided advice on bird diets and donated Mazuri Mini Bird Diet for this study. The Hill and Hood labs at Auburn University provided insightful comments on an earlier version of the manuscript.

## Data availability

All data will be publicly available on Dryad upon manuscript acceptance.

