## Supplemental Tables and Figures for "Captivity affects mitochondrial aerobic respiration and carotenoid metabolism in the house finch (*Haemorhous mexicanus*)"

### SUPPLEMENTARY INFORMATION

**Table S1.** Results of linear models testing for effects of captivity (i.e. caged vs. free-living wild house finches) on mitochondrial respiration and carotenoid concentration measures, including fixed effects of age and molt percent.

| Response variable | Effect | Estimate | Std. Error | *t* | *P* |
|---|---|---|---|---|---|
| Non-ketocarotenoid concentration (post-experiment) | Intercept | 2.90 | 1.22 | 2.38 | 0.02 |
| | Captivity (wild) | 0.05 | 0.35 | 0.14 | 0.89 |
| | Age (hatch year) | -0.15 | 0.33 | -0.45 | 0.65 |
| | Molt percent | -0.01 | 0.01 | -0.41 | 0.68 |
| 3-OH-echinenone concentration (post-experiment) | Intercept | -2.05 | 2.35 | -0.87 | 0.39 |
| | Captivity (wild) | 2.85 | 0.67 | 4.27 | <0.001 |
| | Age (hatch year) | 0.15 | 0.64 | 0.24 | 0.81 |
| | Molt percent | 0.05 | 0.03 | 1.89 | 0.06 |
| Complex I state 3 respiration | Intercept | 6.25 | 0.48 | 12.90 | <0.001 |
| | Captivity (wild) | -0.54 | 0.14 | -3.97 | <0.001 |
| | Age (hatch year) | 0.14 | 0.13 | 1.06 | 0.30 |
| | Molt percent | 0.0037 | 0.0052 | 0.706 | 0.48 |
| Complex I state 4 respiration | Intercept | 0.97 | 0.013 | 75.89 | <0.001 |
| | Captivity (wild) | -0.011 | 0.0036 | -3.03 | 0.0039 |
| | Age (hatch year) | 0.0024 | 0.0035 | 0.69 | 0.50 |
| | Molt percent | 0.00017 | 0.00014 | 1.22 | 0.23 |
| Complex I RCR | Intercept | 16.13 | 4.06 | 3.97 | <0.001 |
| | Captivity (wild) | -1.29 | 1.15 | -1.12 | 0.27 |

| | | | | | |
|---|---|---|---|---|---|
| | Age (hatch year) | 0.52 | 1.12 | 0.47 | 0.64 |
| | Molt percent | -0.041 | 0.045 | -0.912 | 0.37 |
| Complex II state 3 | Intercept | 5.00 | 0.50 | 10.08 | <0.001 |
| | Captivity (wild) | -0.22 | 0.14 | -1.56 | 0.12 |
| | Age (hatch year) | 0.20 | 0.14 | 1.46 | 0.15 |
| | Molt percent | 0.0024 | 0.0054 | 0.44 | 0.66 |
| Complex II state 4 | Intercept | 4.04 | 0.40 | 10.21 | <0.001 |
| | Captivity (wild) | -0.23 | 0.11 | -2.12 | 0.039 |
| | Age (hatch year) | 0.18 | 0.11 | 1.67 | 0.10 |
| | Molt percent | 0.0024 | 0.0043 | 0.56 | 0.58 |
| Complex II RCR | Intercept | 0.30 | 0.064 | 4.64 | <0.001 |
| | Captivity (wild) | 0.041 | 0.017 | 2.40 | 0.021 |
| | Age (hatch year) | 0.015 | 0.017 | 0.89 | 0.38 |
| | Molt percent | 0.00025 | 0.00070 | 0.36 | 0.72 |

When applicable, the reference group for a categorical variable is listed in parentheses.

**Table S2.** Results of a linear model testing the effects of initial (pre-experimental) levels of circulating 3-OH-echinenone, lutein, and zeaxanthin, as well as molt percent, cage size treatment, and age, on final levels of circulating 3-OH-echnineone in the captive birds.

| Effect | Estimate | Std. Error | *t* | *P* |
|---|---|---|---|---|
| Intercept | -1.07 | 1.90 | -0.57 | 0.58 |
| Initial 3-OH-echinenone | 0.10 | 0.01 | 6.48 | <0.001 |
| Initial lutein | -0.15 | 0.12 | -1.30 | 0.20 |
| Initial zeaxanthin | 0.38 | 0.56 | 0.67 | 0.51 |
| Molt percent | 0.02 | 0.02 | 0.84 | 0.41 |
| Cage size (small) | 0.55 | 0.38 | 1.45 | 0.16 |
| Age (hatch year) | -0.34 | 0.51 | -0.67 | 0.51 |

When applicable, the reference group for a categorical variable is listed in parentheses.

**Table S3.** Results of a linear model testing the relationship between final (post-experimental) circulating 3-OH-echinenone and concentrations of 3-OH-echinenone detected in growing feather follicles, with molt percent, cage size, and age as fixed effects.

| Effect | Estimate | Std. Error | *t* | *P* |
|---|---|---|---|---|
| Intercept | 0.45 | 1.87 | 0.24 | 0.82 |
| Circulating 3-OH-echinenone | 0.21 | 0.052 | 4.00 | 0.001 |
| Molt percent | 0.025 | 0.020 | 1.25 | 0.23 |
| Cage size (small) | 0.60 | 0.54 | 1.06 | 0.31 |
| Age (hatch year) | 1.13 | 0.49 | 2.30 | 0.037 |

When applicable, the reference group for a categorical variable is listed in parentheses.

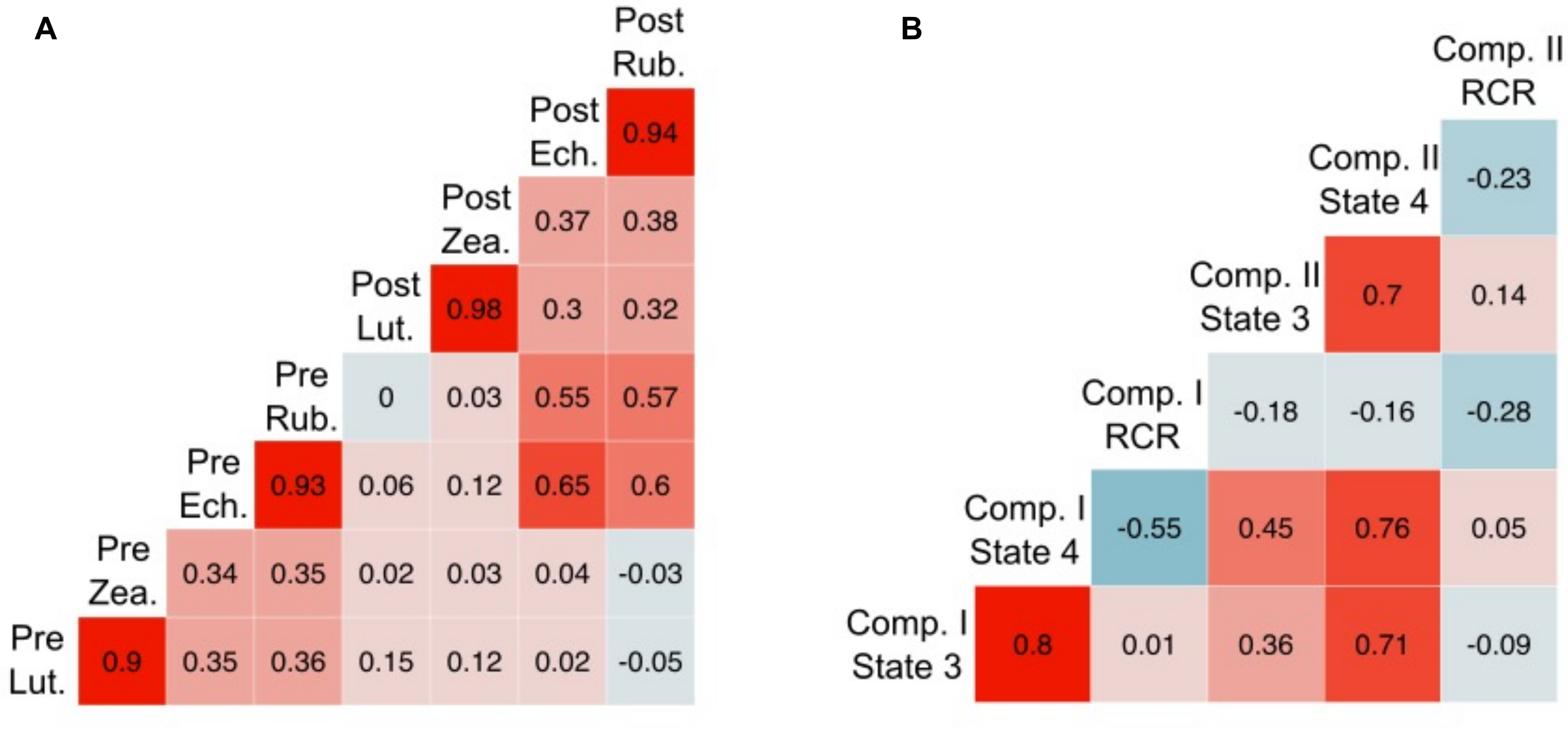


**Figure S1. Pearson correlations among (A) pre- and post-experimental circulating carotenoid concentrations, and (B) mitochondrial respiration measures among captive birds.** Comp. = complex, RCR = respiratory control ratio, Lut. = lutein, Zea. = zeaxanthin, Ech. = 3-OH-echinenone, Rub. = 4-oxo-rubixanthin.

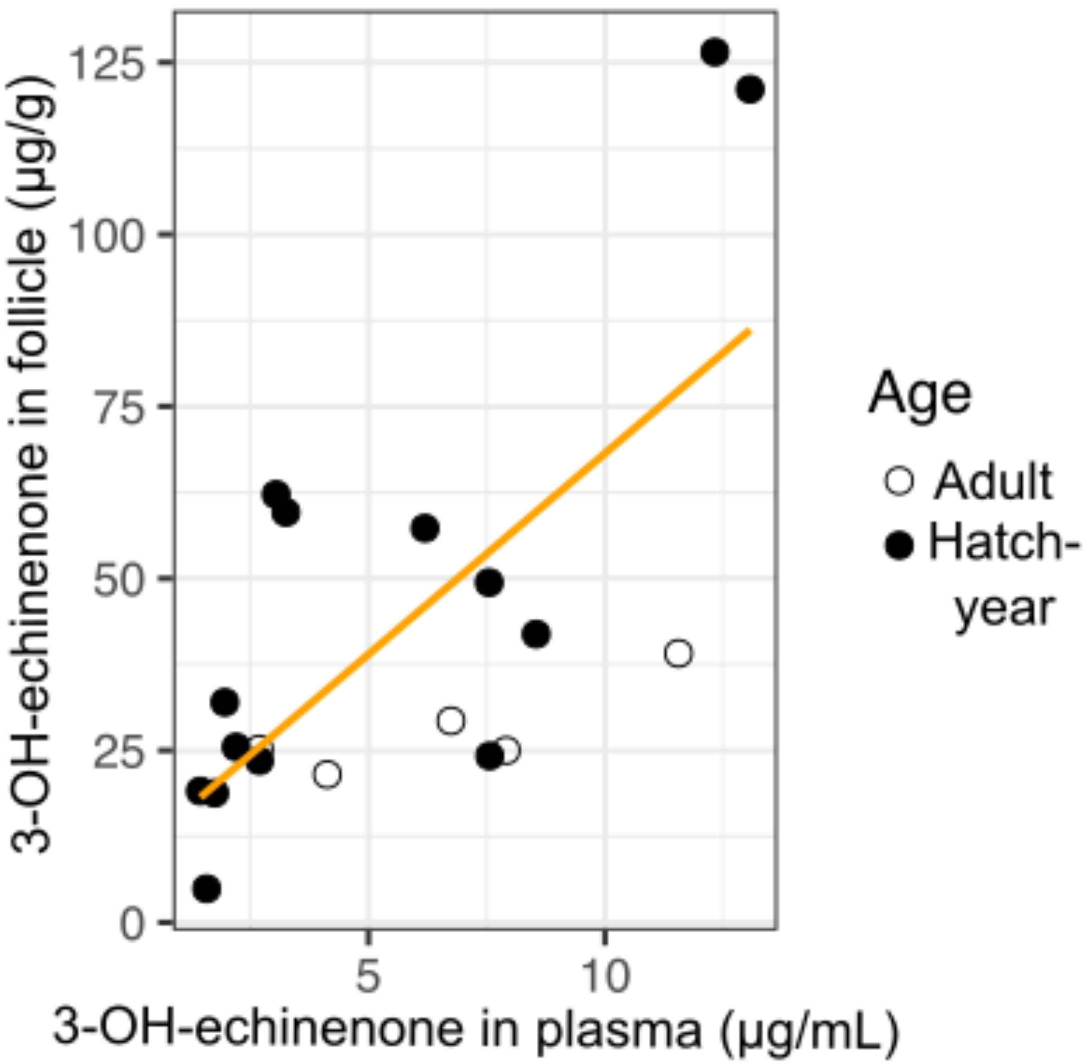


**Figure S2. Scatterplot of concentrations of 3-OH-echinenone measured in circulation *versus* in growing feather follicles of captive-held birds.**
